# Diffusion model of delipidation in biological sample clearing

**DOI:** 10.1101/2023.06.18.545453

**Authors:** Jinglu Han, Xinyi Liu, Xiaoxiao Hou, Yuejia Zhong, Zhiqiang Chen, Zhenyi Yang, Tianzi Jiang

## Abstract

Biological sample clearing techniques are a potent tool for three-dimensional biological imaging, among which delipidation is an essential step in achieving high-quality biological sample transparency. Detergents and organic solvents can both be used for lipids removal. The former has been extensively investigated in biological sample clearing, while the delipidation process based on organic solvents remains to be further elucidated. Recently, organic solvents also served as a delipidation reagent in aqueous-based clearing methods and exhibited very fast clearing speed. To explain the high efficiency of organic solvents, we described the delipidation process of both detergents and organic solvents with a simple diffusion model, we proposed a possible mechanism of the delipidation process of water-miscible polar organic solvents based on the clearing results of brain samples. Both our results and model revealed that polar or non-polar organic solvents with a certain molecular structure could achieve a much faster clearing speed than detergents which could be a guide for establishing a rapid clearing protocol for biological samples with large volumes.

## Introduction

In the past decade, different biological sample clearing methods have been developed to satisfy various volumetric imaging requirements^1^. All these clearing methods can be roughly classified into two groups: one is the hydrophilic method, and another is the hydrophobic clearing method. Lipids are strong light scattering substances thus delipidation is essential for high-quality clearing^1^. Traditionally, hydrophilic methods like CLARITY^2^ or CUBIC^3^ use detergents such as SDS, and Triton X-100 for delipidation. However, the detailed delipidation process is still not very clear in organic solvents based(“solvents-based” in brief) clearing method. Previous studies show that lipophilic dyes are not compatible with hydrophobic methods which implies that lipids have been removed during their clearing process^4^. Water-miscible organic solvents(methanol, ethanol, tert-butanol, THF, etc.) may be served as both delipidation and dehydration reagents while water-immiscible organic solvents such as DCM can be only used for further delipidation in hydrophobic clearing methods^5^. For example, iDISCO^6^, BABB^7^, 3DISCO^5^, and uDISCO^8^ protocols use methanol, ethanol, THF, and tert-butanol(TBA) for dehydration and delipidation may also occur during dehydration while DCM was only used as a delipidation reagent. Recently, only utilizing the delipidation property of THF and combining hydrophilic refractive index matching(RI), Fast3D^9^ and EZ Clear^10^ were developed as more rapid aqueous clearing protocols compared with traditional detergent-based aqueous methods. These results imply that organic solvents such as THF have powerful delipidation abilities and are likely faster than detergents-based clearing methods.

Here we aimed at clarifying the delipidation process based on organic solvents and why organic solvents with certain structures could realize faster clearing speed than a detergent-based solution. Indeed, our results demonstrated that organic solvents exhibit much faster delipidation speed than detergent-based recipes and even faster than CHAPS-based clearing^11^ reagent which once helped penetrate and clear the whole human brain. To decipher the rapid delipidation process of organic solvents, we used a diffusion model to describe the delipidation process of both organic solvents and detergents based on our brain sample clearing results, then we presented a new model to describe the delipidation process of water-miscible organic solvents. We hypothesized that the size of lipids transporting particles in organic solvents is much smaller than that of detergents-based delipidation reagents which may be the most important factor that affects efficiency. A much smaller transporting particle is less hampered by complex 3D biological sample structure and thus is easily washed out from the inside of biological samples, especially for the clearing of large samples. The size of lipid transporting particles combined with other factors like viscosity, biological sample thickness changes, and lipids proportion of transporting particles may jointly explain the delipidation efficiency difference between detergents and organic solvents.

In summary, our results and diffusion model give a clue that organic solvents with certain structures such as THF and DCM are much faster delipidation reagents compared with detergent-based solutions and are good options for establishing rapid hydrophilic clearing protocol or clearing methods for large biological samples or organ.

## Results

Delipidation-based biological sample clearing methods tend to achieve good biological sample transparency as lipids are strong light-scattering substances. Delipidation and RI matching are the two main steps in these methods. Detergents and organic solvents can both be used for delipidation, here we focused on investigating the delipidation process and the mechanism of organic solvents. The degree of biological sample transparency was often affected by the degree of delipidation. Thus the final biological samples transparency can be served as an indicator for the degree of delipidation if only two criteria are satisfied: one is that no delipidation occurs in the RI matching step, and another is that the RI matching solution itself could not make undelipidated biological samples significant transparent. We first eliminated solvents-based RI matching liquid in which delipidation may also occur in this step. To make biological samples transparent, solutions with certain RI value is essential, among various RI matching solution, iohexol-based cocktails (RIMS^12^, EasyIndex^13^ eg.), antipyrine based cocktail(CUBIC-R^14^ eg.), m-Xylylenediamine(MXDA^15^) based cocktail(MACS-R2^15^ eg.) were our candidates, however, iohexol is too expensive to test, MACS-R2 tend to make undelipidated biological samples transparent due to its powerful hyperhydration ability. After confirming that CUBIC-R(+)^14^ could not make undelipidated biological samples transparent(Figure 1B), we adopted CUBIC-R(+) as RI matching solution of biological samples delipidated with organic solvents or detergents.

**Figure 1.**
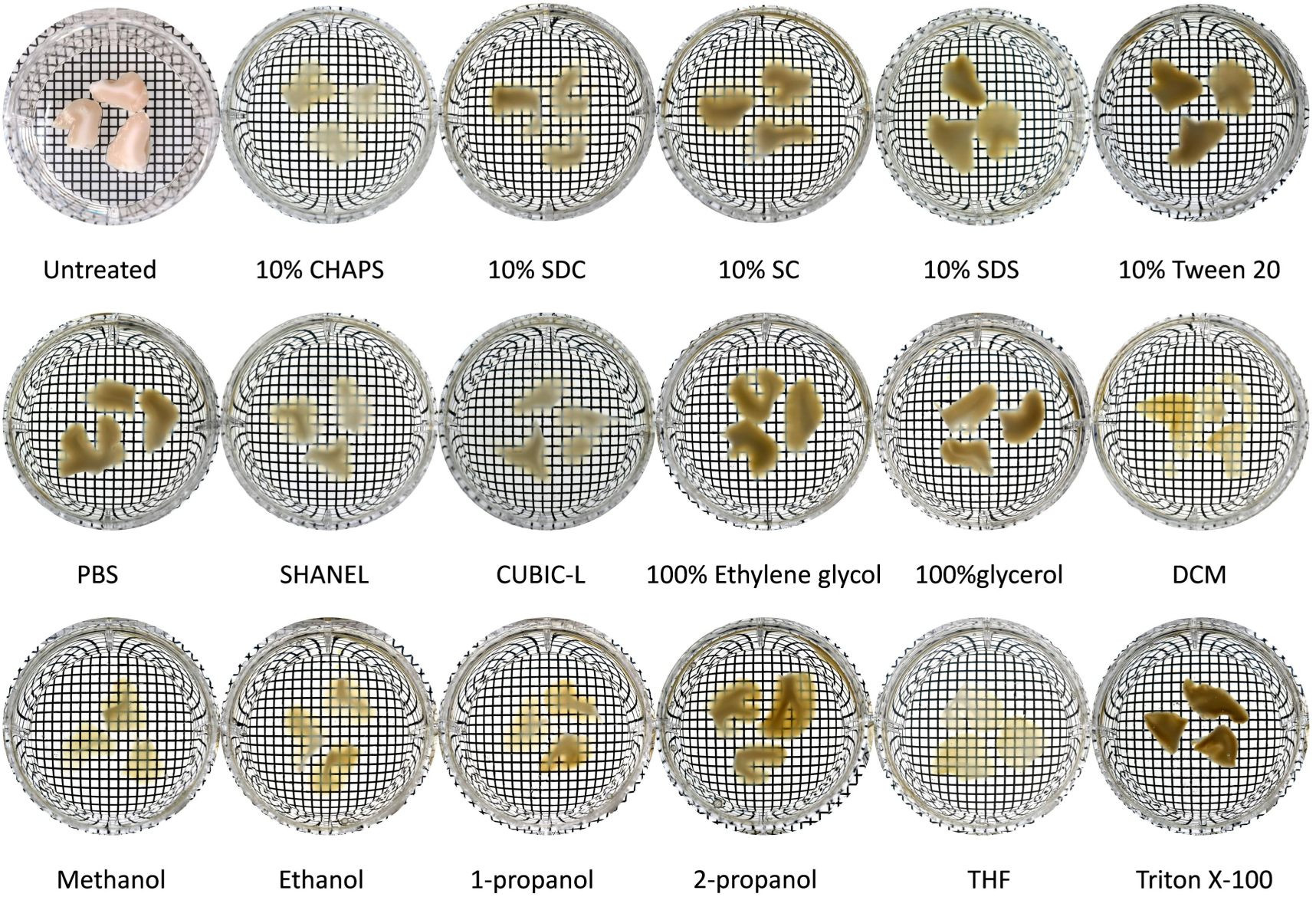
Delipidation efficiency of detergents and various organic solvents.

### The delipidation ability of most of our tested organic solvents was better than detergent-based hydrophilic clearing solution

To investigate the delipidation mechanism of organic solvents, we compared the clearing ability of commonly used aqueous detergent-based delipidation reagents with organic solvents under the same condition(Figure 1). We found that apart from 2-propanol, nearly all tested organic lipid-extracting solvents outperformed detergent-based solutions, especially for the clearance of brain white matter even CHAPS and its analogs like SC, SDC-based recipes were less efficient(Figure 1).

Among all tested solvents, water-miscible solvent THF and water-immiscible solvent DCM successfully achieved the clearing of both brain gray and white matter which exhibited much more powerful delipidation ability compared with other solvents(Figure 1). It seems that THF was slightly better than DCM for white matter clearing(Figure 1). However, the clearing ability between THF and DCM still needed to be further tested.

Apart from the above organic solvents, we also tested the delipidation ability of pure ethylene glycol or glycerol, as we expected, they are not efficient(Figure 1).

### An explanation of the delipidation process of both detergents and organic solvents delipidation using a diffusion model

Why delipidation using organic solvents was much faster than that of detergents?

We assumed three stages of the delipidation process of lipid-extracting solution (solvents): dissolution, forming the micelle(optional), and diffusion. The lipids within biological samples first dissolve by solution(solvents), then lipids may aggregate into micelles, and finally, lipids diffuse out of biological samples. We considered that the diffusion stage of delipidation also conformed to Fick’s law(equation[1]) which has been used for describing 3D antibody staining^16-18^.

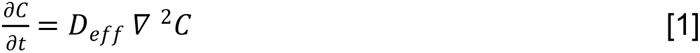

Where:

*∂C*/*∂t* represents the rate of change of the solute concentration with respect to time.

*D*_*eff*_ is the effective diffusion coefficient, which characterizes how readily the solute diffuses in a particular medium.

∇^2^*C* is the Laplacian operator of the solute concentration, representing the gradient of concentration.

In the context of delipidation in biological sample clearing, the solute refers to the lipid molecules or micelles, and the concentration (*C*) represents the lipid concentration within the biological samples.

The equation[1] describes how the lipid-extracting solution (solvent) diffuses into the biological samples and interacts with the lipids, causing them to dissolve and diffuse out. The rate at which this diffusion process occurs depends on the diffusion coefficient of the solvent in the biological samples and the concentration gradient of lipids within the biological samples.

Fick’s second law of diffusion and the Stokes-Einstein equation are related through the diffusion coefficient (*D*) and the properties of the diffusing particles. Fick’s second law of diffusion, as mentioned earlier, describes the rate of change of solute concentration over time in terms of the diffusion coefficient and the concentration gradient. It provides a macroscopic description of diffusion in a system. The Stokes-Einstein equation is a fundamental relationship that relates the diffusion coefficient of a particle to its size and the properties of the surrounding medium.

Ideally, diffusion constant(*D*_0_) can be described by the Stokes-Einstein equation which is given below equation:

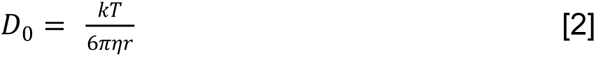

Where:

*D*_0_is the diffusion coefficient of the particle.

*k* is Boltzmann’s constant.

*T* is the temperature.

*η* is the dynamic viscosity of the medium.

*r* is the radius of the diffusing particle.

The equation suggests that the diffusion coefficient is inversely proportional to the size of the particle (radius) and directly proportional to the temperature and the reciprocal of the medium’s viscosity.

The relationship between Fick’s second law and the Stokes-Einstein equation arises when considering diffusion in a system where the diffusing particles are affected by both concentration gradients and the properties of the surrounding medium. In the case of delipidation in biological samples clearing, the lipid-extracting solution (solvent) acts as the medium, and the lipids within the biological samples act as the diffusing particles. The equation[2] described the case of the diffusion constant in pure solution(solvent). However, lipids transporting particles are not only suffered by the surrounding medium, but also by the biological gel mesh, especially for the large particles. Thus the effective diffusion constant(D_*eff*_) of lipids should be corrected under different conditions. Senanayake et al. summarized the scaling theory of contrast size-dependent diffusion of nanoparticles within entangled solutions versus cross-linked gels^19-21^. The theory gives a calculating model of the modification of particles in different particle size ranges^19-21^. For example, if the particle size is far smaller than the correlation length(*ξ*), the particles experience mostly the solvent viscosity while for particles slightly larger than the mesh size (*a*_*x*_) the diffusion coefficient D ∼ *exp*[−(2r/*a*_*x*_)^2^]. *r* is the particle radius, *a*_*x*_ is the average distance between two permanent cross-links^19-21^.

In all, lipid diffusion is hindered by biological gel structure which decreases the diffusion process. Thus diffusion constant needs to be corrected by considering the particle size of lipids, the degree of fixation of biological samples, and so on. For simple, we utilized the effective diffusion coefficient(*D*_*eff*_) to represent the real coefficient in Fick’s law(equation[1]) though we did not know how the effective coefficient was precisely calculated.

To explain the rapidness of solvent-based delipidation, we discussed the parameters in the above equations.

A simple relation between delipidation time(*t*) and sample thickness(*d*), effective diffusion constant(*D*_*eff*_)can be written as below according to equation[1]:

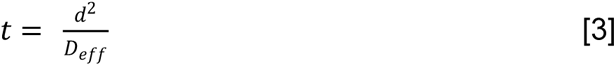

For the given biological sample clearing method, raising the temperature(*T*) and thinner biological sample thickness(*d*) are strategies for accelerating delipidation speed and shortening clearing time according to the above equation[2]. For example, Yu et al.^22^ and Murray et al. successfully sped the clearing time of PACT^12^ and SWITCH^23^ by elevating the temperature. SHIELD^24^ shortens clearing time by dissecting the whole mouse brain into hemispheres and it’s obvious that brain slices are easier to be cleared compared with the whole brain.

To compare the delipidation speed difference between organic solvents and detergent-based solution, we fixed parameters *d* (using brain slices with the same thickness) and *T*(performing tests under the same temperature), so the difference in delipidation efficiency between organic solvents and detergents lies in parameter viscosity(*η*) and particle size(*r*) according to above equation[2][3].

We first discussed the parameter viscosity((*η*), and considered that the viscosity lipid suffered during diffusion is affected by the resistance of the solution(solvents), the biological samples’ gel mesh structure, and the volume of particles. Diffusion coefficient is reciprocal to the medium’s viscosity((*η*). Among our tested organic solvents, THF and DCM were much more efficient than others(Figure 1), low viscosity may also improve their efficiency as no intermolecular hydrogen bonds are formed like alcohols. We also tested 100% Triton X-100 for delipidation(Figure 1), as we expected, they are all not efficient, the less efficiency may be explained by its high viscosity. Apart from the viscosity of the delipidation solvents or solution, lipid transporting particles are also restrained by the biological gel mesh structure(delipidated biological samples can be regarded as electrolyte-gel^16^) in which lipids are subjected to greater viscous resistance during transportation within the biological samples to some extent, we regard this as an inner viscosity. The inner viscosity from biological gel mesh exhibits a decrease in the diffusion constant which may be corrected by the scaling theory. For example, the inner viscosity increases in biological samples delipidated by ionic detergent(SDS eg.) or organic solvents due to the shrinkage of biological samples. On the contrary, inner viscosity decreased in biological samples delipidated with some low-concentration of non-ionic detergents(like Triton X-100) due to the swelling of biological samples, this assumption may explain why Triton X-100 based protocol(CUBIC-L) are 3-7 days while SDS based clearing needs up to 7 days for whole mouse brain clearing according to previous reports^3,25^. However, the clearing speed between SDS or Triton X-100 at low solutes concentration(around 10w/v) still needs experiments to be compared.

Next, we focused on the parameter *r* (particle size), which represents the particle radius that transports lipids. Traditionally, hydrophilic methods(CLARITY, CUBIC eg.) utilized detergents like SDS or Triton X-100 for delipidation, in which lipids are transported in the way of micelles from the inside of biological samples(Figure 2B, 2C). However, large micelles of these detergents reduced the diffusion efficiency, especially for clearing large animal organs.

**Figure 2.**
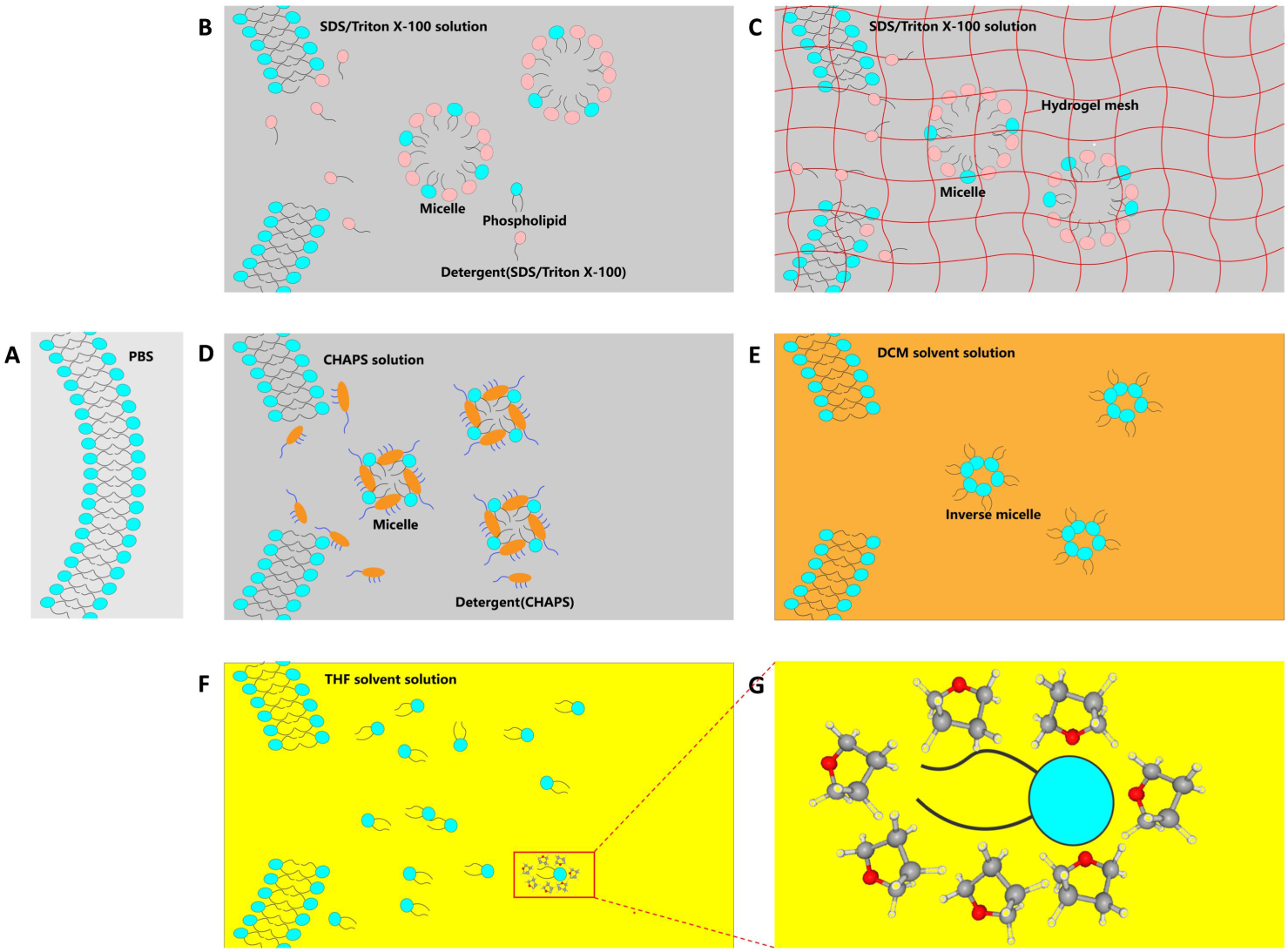
Diffusion model of lipid removal. **(A)**Intact cell membrane without delipidation (membrane proteins, glycoproteins, or other components are not shown here for simple) **(B)**Large lipid transporting particles (micelles) are formed using detergents(SDS/Triton X-100) as long as their concentration reaches critical micelle concentration(CMC) due to the typical “head-to-tail amphipathic structure of these detergents. Large micelles increase diffusion viscosity thus reducing clearing efficiency. **(C)**Similar model described in Figure 2B, the difference is that biological samples is just pre-embedded with hydrogel in which delipidation efficiency may be further reduced due to the resistance of cross-linked gel. **(D)**Detergents like CHAPS contain a hydrophilic tail and atypical facial amphipathic including a rigid steroidal structure with a hydrophobic convex side, and a hydrophilic concave side (bearing three hydroxyl groups). With the special hydrophilic tail and amphipathic head, CHAPS has higher CMC and forms smaller micelles compared with SDS and Triton X-100, which are more permeable in large biological samples or organs. **(E)**Phospholipids are also amphipathic which automatically form inverse micelles^26^ in a hydrophobic environment(DCM eg.) to hide their hydrophilic head for maintaining the stable status of the whole system. Here we hypothesized that the size of inverse micelles is also smaller than the micelles formed by SDS/Triton X-100, even smaller than CHAPS as the size proportion of the hydrophilic head of phospholipids is smaller than that of the hydrophobic tail. **(F)**We assumed a new model that phospholipids diffused freely within polar hydrophilic organic solvents(THF eg.) as both phospholipids and surrounding organic solvents are amphipathic thus no micelles are formed. **(G)**Magnified view of a simplified model of the arrangement of THF molecules surrounding phospholipid. Phospholipid does not need to hide their head or tail thus no micelles are formed within amphipathic solvents.

To solve this problem, Zhao et al. introduced CHAPS in their SHANEL^11^ method in which CHAPS formed small micelles compared with SDS or Triton X-100 due to the special structure of CHAPS containing a hydrophilic tail and an atypical amphipathic head with a rigid steroidal hydrophobic convex side and a hydrophilic concave side bearing three hydrophilic hydroxyl groups(Figure 2D). Due to the high permeability of small micelles, SHANEL successfully achieved whole human or large animal organ clearing.

### A new lipids diffusion model of water-miscible organic solvents

As for solvents-based methods, lipids transporting particles may differ from that of detergents, Richardson et al.^26^ summarized four ways of delipidation including both detergents and organic solvents, and they came up with the concept of the inverse micelle(Figure 2E) formed in hydrophobic liquid DCM which is a commonly used delipidation solvent in 3DISCO, uDISICO, SHANEL, etc^26^. Inverse micelles of phospholipids are automatically formed to hide their hydrophilic heads in the hydrophobic environment of DCM to maintain the lowest system energy due to the amphiphilic structure of phospholipids(Figure 2E).

However, the inverse micelles model could not explain the delipidation process of the polar hydrophilic organic solvents (THF and low alcohol such as methanol, ethanol, etc.) as whose structure differs a lot from non-polar hydrophobic solvents. Here we assumed a new model of the delipidation process of water-miscible organic solvents, we hypothesized that no micelles or only very small lipid clusters are formed and the lipids are transported in the way of a single molecule or small clusters because water-miscible organic solvents are also amphiphilic in which phospholipids can be dissolved directly without forming micelles(Figure 2F, 2G).

In short, we summarized five types of lipid transporting particles(Figure 2): (1)large micelles in SDS/Triton X-100 clearing, (2)large micelles in SDS/Triton X-100 clearing in hydrogel-embedded samples, (3)small micelles in CHAPS clearing, (4)inverse micelles in DCM clearing, (5)single or small lipid clusters in THF clearing. Our results demonstrated that both DCM and THF are more powerful than detergents(Figure 1A, 1B). It’s easy to explain THF is more powerful than CHAPS as the former forms no micelle. However, both DCM and CHAPS solution forms micelles, so the new question is which micelles are smaller, micelles of CHAPS or inverse micelles of DCM? To answer this question, we need experiments to measure the real size of these micelles. Here we assumed that the inverse micelles of DCM are smaller than that of CHAPS as the size of the hydrophilic head of phospholipids is smaller than that of the tail, thus a small number of lipids are enough to hide their hydrophilic heads in DCM. If this assumption is correct, the sequence of particle size is (1)=(2)>(3)>(4)>(5) under the premise that hydrogel does not change the micelle size.

To systematically answer why organic solvents are much more efficient in lipid removal compared with detergents, we considered the influence of lipid-dissolution efficiency, the effective diffusion constant in Fick’s law, and the lipid transporting efficiency of transporting particles, finally at least four possible reasons were summarized:

1. The dissolution efficiency of solvents is higher than that of detergents solution due to the higher collision probability between lipid molecules and solvents since solvent-based delipidation reagent contains 100% solvents while detergent-based reagent contains only 0-20% delipidation solutes.
2. The lipid transporting particle size(*r*) in organic solvents is much smaller than that of detergents which may dramatically affect the effective diffusion coefficient according to scaling theory, equation[2], and Stokes-Einstein equation.
3. Though solvents based clearing causes biological sample shrinkage which may increase inner viscosity(*η*), shrinkage also reduces sample thickness(*d*) which reduces the distance of diffusion.
4. The proportion of lipids in transporting particles in organic solvents is much higher than that of detergents since the particles in the detergent-based clearing are composed of both detergent and lipid molecules even lipids only take up a small proportion of the particle which reduce the transporting efficiency while the particles in organic solvents clearing consist of only lipids(Figure 2B-G). The above factors may jointly account for the rapidness of organic solvents.

Based our new assumption and discovery, we have developed various much efficient delipidation strategies under the guidance of diffusion model, which further increase the reliability of the diffusion model(Figure 3, strategies 1-7, or s1-s7).

**Figure 3.**
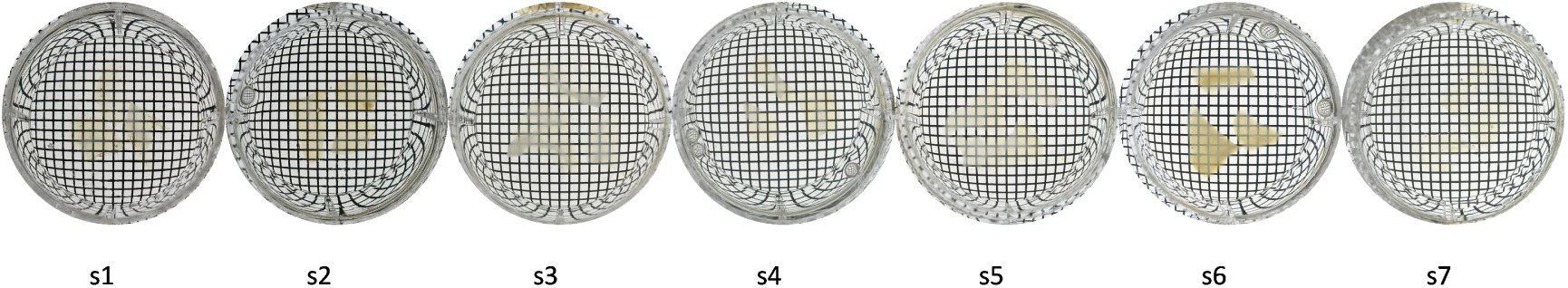
Establishment of various efficient strategies of delipidation under the guidance of diffusion model.

## Discussion

Delipidation is a crucial step for high-quality clearing, however, delipidation process of organic solvents is rarely systematically studied. Our results indicated that organic solvents with certain structures exhibited better delipidation efficiency than detergent reagents even CHAPS could not surpass these organic solvents. In this article, we comprehensively discussed the delipidation process of both hydrophobic and hydrophilic reagents with a diffusion model. We proposed a new diffusion process of water-miscible organic solvents in which no micelles are formed during this process. We considered that the small size of lipids transporting particles contributed a lot to the rapid delipidation of organic solvents, other factors like viscosity(*η*), particle size(*r*), and sample thickness(*d*) jointly explained the rapid delipidation process of organic solvents.

Among our tested solvents, THF and DCM are both sufficient for clearing 3mm thickness pig slice within 12h under room temperature, and THF seems just slightly better for brain white matter clearing, however, their delipidation ability comparison still needs to be further tested. On the one hand, THF may exhibit higher permeability than DCM as the former forms smaller lipid transporting particles size than the latter; on the other hand, hydrophobic DCM possessing higher logP(lipophilic index) is more lipophilic than THF. Thus THF and DCM exhibited similar delipidation ability of 3mm thick pig slices when considering both transporting particle size and lipophilicity. Here we hypothesized that DCM is more powerful to clear relatively thin slices while THF may be more competent for clearing biological samples with larger volumes due to the high permeability. The above discussion is based on clearing the samples rich in amphipathic phospholipids(brains are typical examples). However, DCM may be more efficient for the clearing of biological samples rich in non-polar fat(triglyceride) like adipose biological samples as both fat and DCM are hydrophobic thus no micelles are formed. Indeed these hypothesis needs our further tests of experiments. Currently, our results of the clearing performance of hydrophilic organics like ethylene glycol or glycerol were very poor(Figure 1) which may indicate that lipophilic parameters(like log P eg.) of organic solvents are also very important for lipid removal other than the influence of the parameters in diffusion function as lipophilic solvents are more likely to dissolve lipids which is the prior step before the diffusion of lipids.

In summary, we explained the rapidness of solvents based delipidation with the help of diffusion function which helps us establish rapid protocols for the clearing of biological samples of large volume or intact organs of large animals like pigs, dogs, or macaques due to the high permeability of some organic solvents. Various efficient delipidation strategies that we have developed may also indicate the reliability of the diffusion model(Figure 3). However, endogenous fluorescent signal quenching is a no neglected problem of most organic solvents, among all our tested solvents, TBA has a relatively better fluorescent signal retention ability^27^, thus rapid hydrophilic biological sample clearing method based on TBA is our direction for the next effort.

## Materials and Methods

### Reagents

| Chemicals | CAS | Vendor, No. |
| --- | --- | --- |
| PBS |  | Solarbio, P1003 |
| PFA | 30525-89-4 | Sigma, P6148-1KG |
| Methanol | 67-56-1 | Sigma, 34860-1L-R |
| Ethanol | 64-17-5 | Meryer, M83117-500ML |
| 1-propanol | 71-23-8 | TCI, P0491 |
| 2-propanol | 67-63-0 | TCI, I0163 |
| Tetrahydrofuran(THF) | 109-99-9 | aladdin, T103263-4L |
| Dichloromethane(DCM) | 75-09-2 | Sinopharm Chemical Reagent Co. Ltd., China, 80047392 |
| (Dimethyl sulfoxide)DMSO | 67-68-5 | Solarbio, D8370 |
| Ethylene glycol | 107-21-1 | Macklin, E808737-500ML |
| Glycerol | 56-81-5 | Vetec, V900122-500ML |
| N-butyldiethanolamine | 102-79-4 | TCI, B0725 |
| Triton X-100 | 9002-93-1 | Sigma, T9284-500ML |
| 3-[(3-cholamidopropyl)dimethylammonio]-1-propanesulfonate(CHAPS) | 75621-03-3 | Solarbio, C8390 |
| N-Methyldiethanolamine | 105-59-9 | Innochem, A45320-500ML |
| Sodium Cholate(SC) | 361-09-1 | TCI, C0325 |
| Sodium Deoxycholate(SDC) | 302-95-4 | TCI, C0316 |
| Sodium dodecyl sulfate(SDS) | 151-21-3 | Sigma, L3771-500G |
| Tween 20 | 9005-64-5 | Aladdin, T104863-500ML |
| Antipyrine | 60-80-0 | TCI, D1876 |
| Nicotinamide | 98-92-0 | Solarbio, N8071 |

CUBIC-R(+) contains 45%wt/wt antipyrine, 30%wt/wt nicotinamide, and 25% ddH_2_O with 0.5%v/v N-butyldiethanolamine for pH adjustment. CUBIC-L is a mixture of 10% (wt/wt) N-butyldiethanolamine and 10%(wt/wt) Triton X-100 in ddH_2_O, SHANEL clearing contains 10% w/v CHAPS and 25% w/v N-Methyldiethanolamine.

### Samples

Whole pig brains were bought from a local slaughterhouse, intact brains were fixed by PFA for 24h, then were dissected into big brain blocs, extra 24h incubation of PFA was for further fixation. 3mm(millimeter) brain slices were sectioned by a vibrating slicer (VT 1200S, Leica) and samples were temporally stored under 4°C if not used immediately.

### Biological sample clearing

For pig brain slice clearing, all samples were immersed in testing solutions or solvents at room temperature for 12h with shaking, then the samples were washed three times (15 minutes each) with ddH_2_O. After washing, samples were incubated in half diluted(50%v/v CUBIC-R(+) diluted with ddH_2_O) for 24h then transferred into CUBIC-R(+) for a further 2 days until biological samples became transparent. Transparent brain slices were put into the 6-well plate for image shots. Neglecting the distortion of tested samples, pig brain slices were directly immersed in 100% organic solvents without pre-incubation. The sample delipidated with DCM was washed 15 minutes*3 times by DMSO before delipidation with DCM in which DCM is water-immiscible and could not be used for delipidation without dehydration.

## Author contributions

Jinglu Han proposed the core idea, and designed and performed the experiments. Xinyi Liu prepared part of the samples, and Jinglu Han, Xiaoxiao Hou, Zhengyi Yang, and Xinyi Liu, Zhiqiang Chen deeply discussed the design of the study and gave useful suggestions, Zhengyi Yang, Yuejia Zhong, Jinglu Han performed the diffusion mathematical modeling. Zhiqiang Chen gave import explanation of the rapidness of the delipidation of organic solvents. Jinglu Han drew the illustration, and Jinglu Han and Zhengyi Yang wrote the manuscript. Tianzi Jiang led and supervised the project. All authors participated in the discussion of the manuscript.

